# Thousands of missing variants in the UK BioBank are recoverable by genome realignment

**DOI:** 10.1101/868570

**Authors:** Tongqiu Jia, Brenton Munson, Hana Lango Allen, Trey Ideker, Amit R. Majithia

**Affiliations:** Department of Medicine, University of California San Diego, La Jolla, CA 92093, USA; Medical Research Council Epidemiology Unit, University of Cambridge School of Clinical Medicine, Cambridge Biomedical Campus, Cambridge, UK

## Abstract

The UK Biobank is an unprecedented resource for human disease research. In March 2019, 49,997 exomes were made publicly available to investigators. Here we note that thousands of variant calls are unexpectedly absent from the current dataset, with 641 genes showing zero variation. We show that the reason for this was an erroneous read alignment to the GRCh38 reference. The missing variants can be recovered by modifying read alignment parameters to correctly handle the expanded set of contigs available in the human genome reference.

## Main Text

The UK Biobank (UKB) is a resource of unprecedented size, scope and openness, making available to researchers deep genetic and phenotypic data from approximately half a million individuals^1^. The genetic data released thus far include array-based genotypes on 488,000 individuals and exome sequencing on 49,997 of these, with further exome sequences to be released in 2020. Such comprehensive cataloging of protein-coding variation across the entire allele frequency spectrum, attached to extensive clinical phenotyping, has the potential to accelerate biomedical discovery, as evidenced by recent successes with other exome biobanks^2^. Given the scale of the data (the current exomes release contains approximately 120 TB of aligned sequence), few investigators have the computational infrastructure or knowledge to identify and curate genetic variants and instead rely on releases of accompanying pre-processed variants in variant call format (VCF, approximately 5 GB). Specifically the UKB has released pre-processed VCFs from two different variant analyses, called the Regeneron Seal Point Balinese (SPB)^3^ and Functionally Equivalent (FE)^4^ pipelines. Although these pipelines are still evolving, studies have already made use of the released exome variants mainly for comparison with previous UKB genotyping data or variant databases^5^. However, a recent report pointed out an error in duplicate read marking in the SPB pipeline that could lead to false variant calls here (http://www.ukbiobank.ac.uk/wp-content/uploads/2019/08/UKB-50k-Exome-Sequencing-Data-Release-July-2019-FAQs.pdf), resulting in the removal of the SPB release from the UKB data repository. Thus, the FE pipeline is currently the only source of variant calls available for downstream research. Here we identify an error in the FE pipeline that results in a systematic lack of variant calls for thousands of genes, along with a solution to patch this bug.

In our initial investigations of protein-coding variation in the UKB exomes, we noted a complete absence of variation in a number of genes of interest, including *CLIC1, HRAS, TNF*, and *MYH11* (one of the ACMG 59 genes in which incidental sequencing findings should be reported^6^). Such absence was unexpected given the UKB exome sample size, as these genes were not under severe evolutionary constraint^7^, and protein-coding variants had been called for these genes in other databases^8^, some of which were present at sufficiently high frequency to be included on genotyping arrays. We reasoned that the lack of variant calls in these genes was unlikely to be explained by ascertainment of a unique population in the UKB (i.e. the variants truly did not exist), and was instead caused by a technical error in sequencing, data processing, variant calling or a combination of those.

In order to prove that the missing variants are indeed present in the UKB population, we first evaluated the internal consistency between the genotyping and exome sequencing data that had been collected for the same UKB samples. In particular, we identified a total of 30,979 common variants (MAF > 0.01) in the UKB dataset that had been ascertained in 49,909 samples by genotyping arrays (**Online Methods**). While the majority of variants had been called by both methods (24,614 variants, 79.5%) a substantial minority (6,365 variants, 20.5%) were called by the genotyping arrays but not by exome sequencing (**Fig. 1**). This discrepancy included many common variants with MAFs close to 0.5 (i.e. that were present in almost 50% of the array samples) providing strong evidence that the exome sequencing genotype calls are lacking variants that exist and should have been detected in this UKB exome population.

**Fig. 1:**
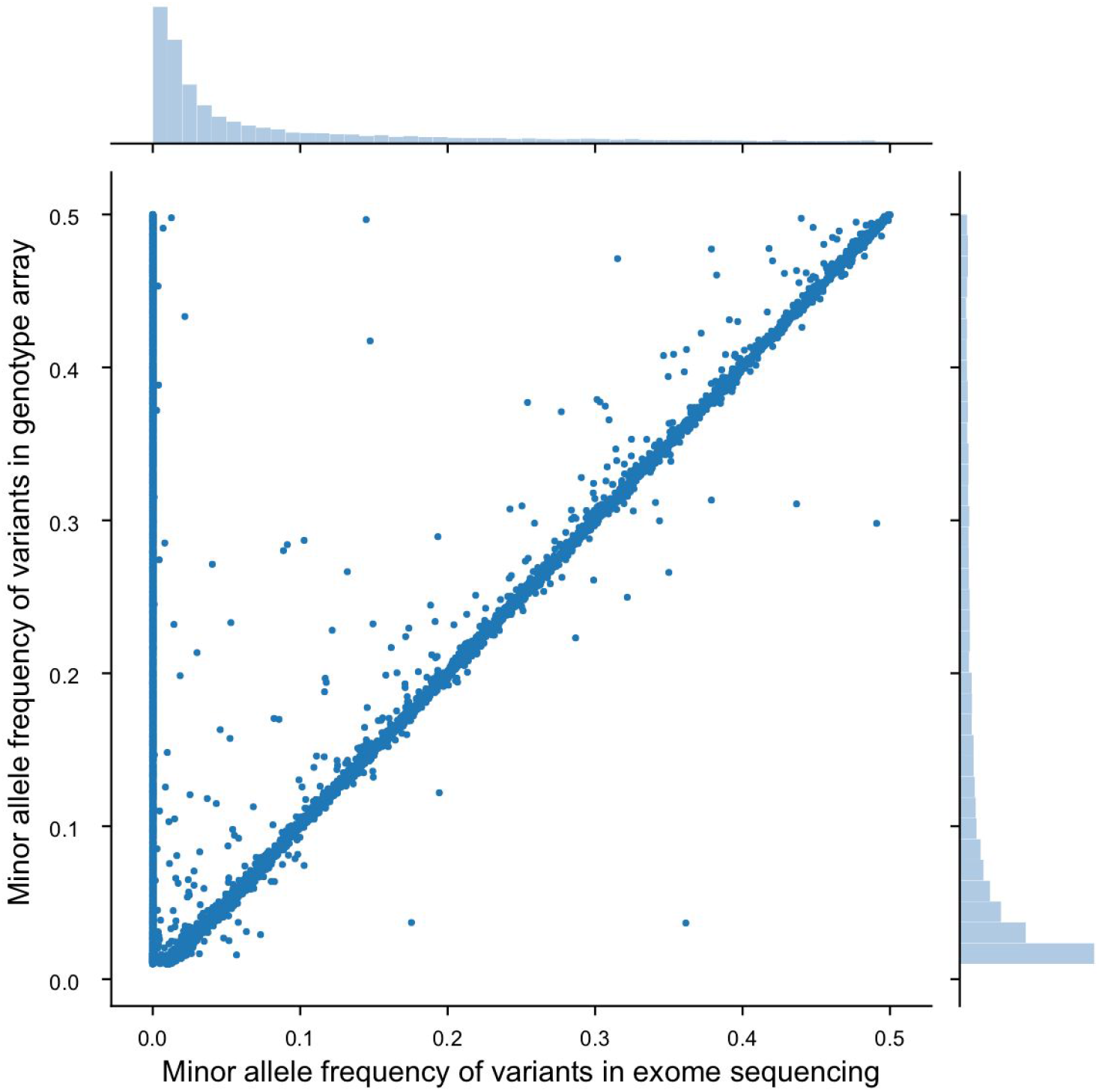
Variant allele frequencies for individuals in the UK Biobank called by analysis of genotyping arrays versus exome sequencing. The Minor Allele Frequency (MAF) determined by each method is plotted, covering a total of 30,979 common variants measured by both methods over 49,909 individual samples. The distribution of variant allele frequencies is shown for each method by histograms above (exome) and to the right of (genotyping array) the main scatterplot.

We next examined variant calls aggregated per gene in the UKB exomes in comparison to the Genome Aggregation Database^9^ (gnomADv2.1.1, 125,748 sequenced exomes, **Online Methods**). Our analysis focused on the exons sequenced both in UKB and gnomAD, which encompasses 23,040 human genes (**Online Methods; Fig. 2a**). We found that, for most genes, the number of variants in gnomAD was well predicted by the number in UKB, with expected 1:2.3 proportionality given the larger gnomAD sample size (**Fig. 2b**). However, this analysis also highlighted 641 genes with 0 variants called in the UKB exomes, versus a median of 286 variants (range of 1 to 14,291) in gnomAD (**Supplementary Table 1**). Using the aggregate observed variant frequency per gene in gnomAD, we calculated the probability for at least one variant being observed in the UKB exome sample for each gene. Of the 641 genes, 598 (93%) should have had at least one variant identified (95% CI one-tailed binomial distribution). Given that the UKB is a predominantly European ancestry population and the gnomAD dataset contains a more diverse population, we performed ancestry-specific analysis (**Fig. 2c**) of these genes in gnomAD. The largest number of variants in these genes were found in the European ancestry samples as expected by their majority representation in the gnomAD dataset. This excluded the possibility that some or all of the genes lacking variation in the UKB was due to ancestry-specific variation.

**Fig. 2:**
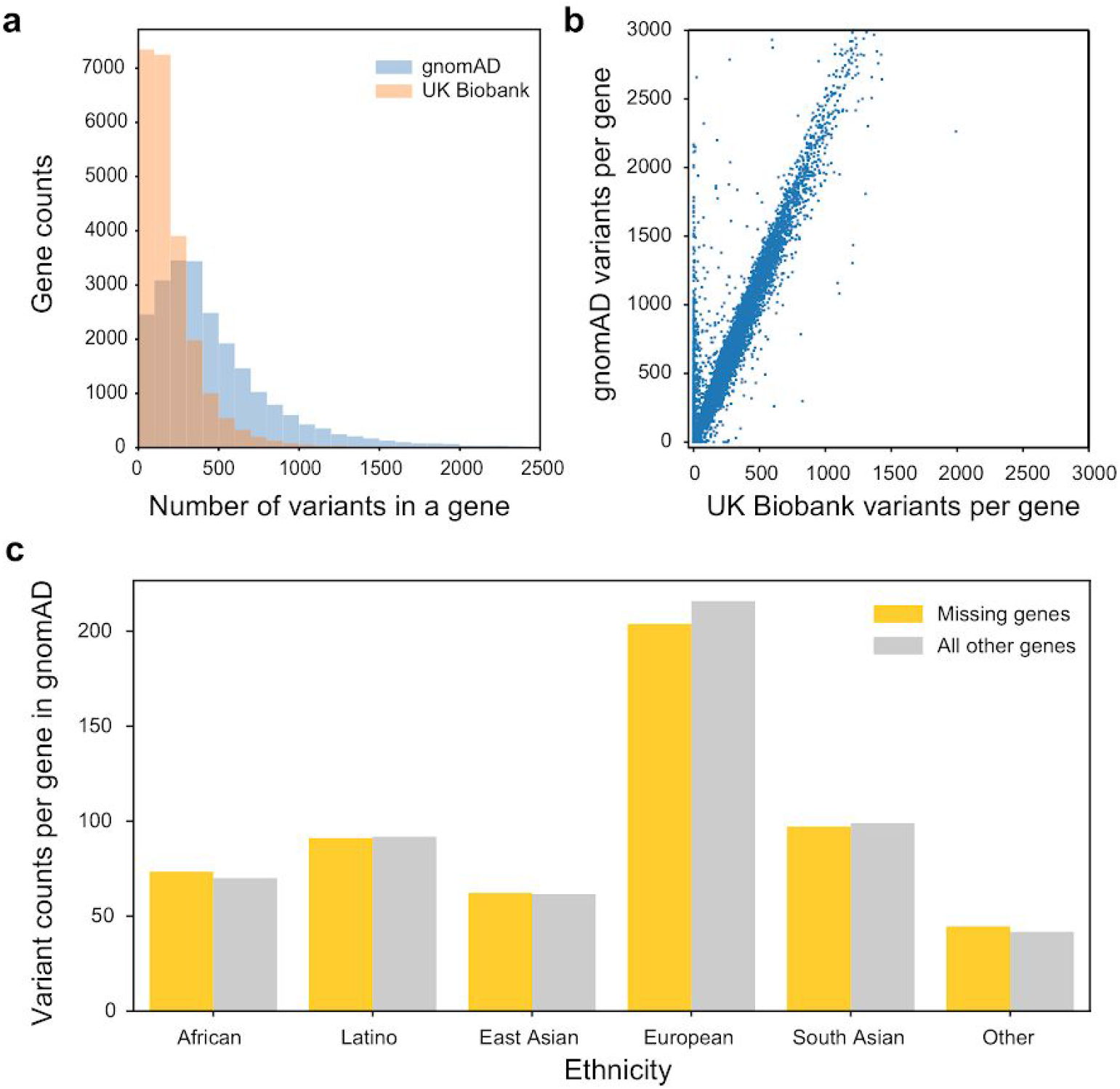
Evaluation of exome variants called by the UK Biobank against the Genome Aggregation Database (gnomAD). **a**, Histogram of variant counts for each of the 23,040 human genes commonly annotated in UKBioBank (orange) and gnomAD (blue), at a fixed bin size of 100. **b**, Scatterplot of variant counts for each gene in gnomAD versus UKBioBank. **c**, Counts for variants in the 641 genes that are covered by gnomAD but missing from UK Biobank (yellow), divided into six subpopulations by ethnicity. Counts for all other human genes are shown as a reference (gray).

To understand the reason for these missing variant calls in the UKB, we analyzed the sequencing read data, provided by the FE pipeline, for individual exomes at each of the 641 loci. Our analysis indicated that, despite having reads mapped to these genes (**Fig. 3a**), the mapping quality (MAPQ) score was universally zero, causing these reads to be eliminated from the downstream procedures for variant calling. The MAPQ field in the SAM specification^10^ is the PHRED scaled probability^11^ of the alignement being erroneous. In practice, however, each aligner treats the MAPQ field differently. With the aligner BWA-MEM^12^ used in FE pipeline, a MAPQ score of zero is given to reads that align equally well to more than one genomic location, and is typically an indicator of reads that come from duplicated or repetitive regions of genomic DNA. However, many of the loci we individually examined were not known to harbor repetitive elements or reside in regions of genome duplication. Investigating further, we found that the zero MAPQ score was due to the reads showing multiple alignments to the GRCh38 genome reference, not to repetitive elements but to so-called ‘alternative contigs’ in the GRCH38 reference (ftp://ftp-trace.ncbi.nlm.nih.gov/1000genomes/ftp/technical/reference/GRCh38_reference_genome/GRCh38_full_analysis_set_plus_decoy_hla.fa). As of this genome release, alternative contigs are used frequently to represent divergent haplotypes which cannot be easily captured by a single linear sequence. Indeed, of the 598 genes with high probability of missing variation, 568 (95%) had alternative contigs represented in the genome reference (**Supplementary Table 1**).

**Fig. 3:**
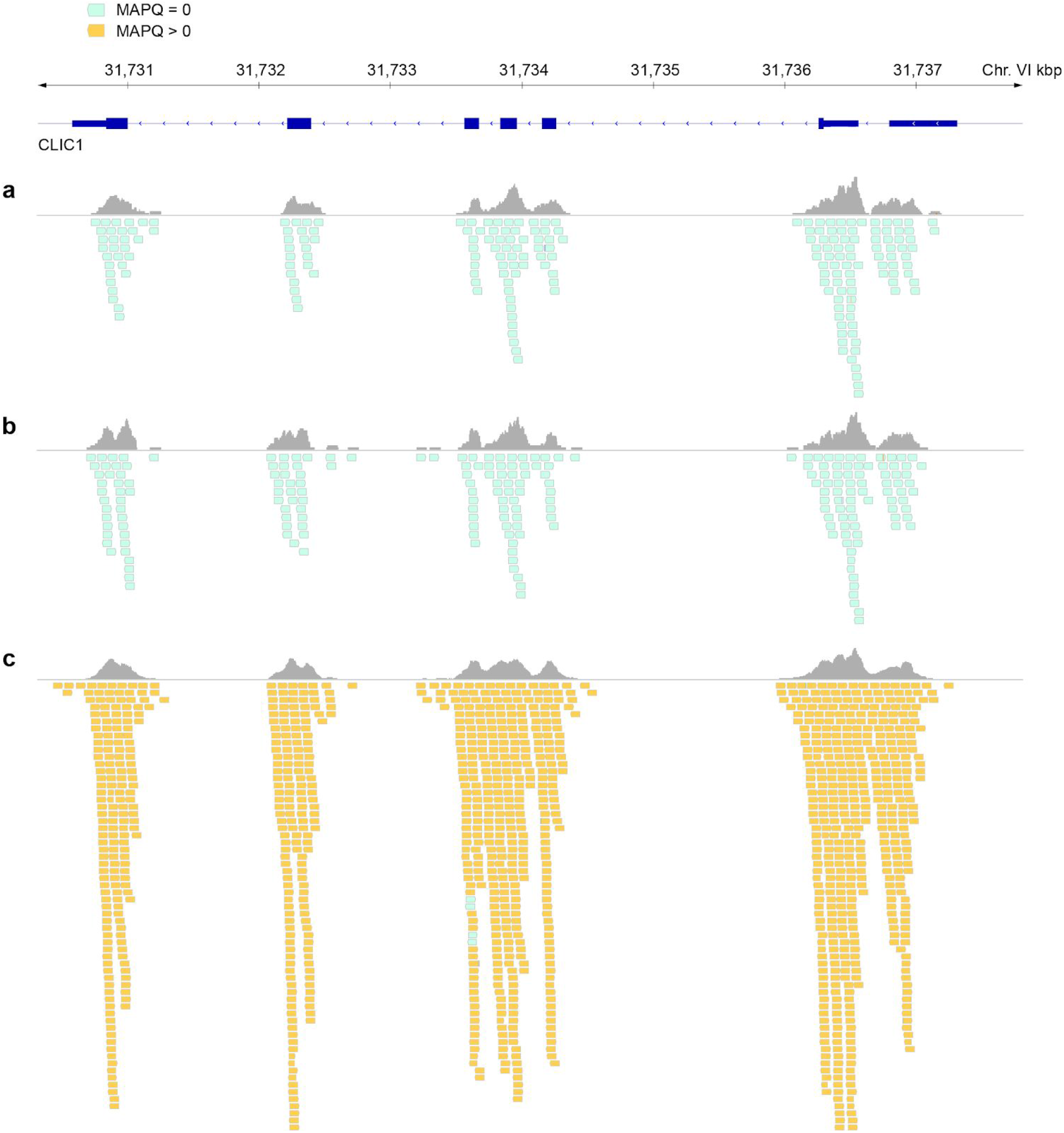
UK Biobank exome read alignments at the CLIC1 genomic locus. **a**, Current alignments obtained from UK Biobank. **b**, Realignment of reads without handling alternative contigs. **c**, Corrected alignments with proper indication of alternative contigs. Read alignments to the human genome are visualized with Integrated Genomics Viewer (GRCh38, IGV version 2.6.2).

Starting from the raw reads available from CRAM files, we found that the original read alignment provided by the UK Biobank (**Fig. 3a**) was most closely reproduced when performing the alignment under default alignment parameters (BWA-MEM^12^, **Online Methods**). This alignment (**Fig. 3b**) does not take into consideration alternative contigs in the absence of an index file specifically marking these contigs, and treats them instead as independent genomic regions equal to primary contigs. Reads that map to both primary and alternative contigs are therefore interpreted as mapping to multiple genomic locations at these loci. We found that re-aligning the raw reads while providing the alternative contig index file for the genome reference resulted in a dramatic improvement in the number of reads that properly mapped to a single genomic locus and therefore had the MAPQ score greater than zero (**Fig. 3c**).

In summary, we have found that genetic variants documented in the UK Biobank FE release are conspicuously absent from certain genes, in a manner that is best explained by errors of read alignment. Furthermore, while our analysis has focused on 641 genes with an absolute lack of variant calls, additional genes may have partially duplicated or repetitive sequences such that they are missing substantial (but above zero) variation beyond those identified in our short study (2391 genes are currently contained within alternative contig representations of the genome). Thus, the variant calls in the current UKB exome data should not be used for large-scale genomic analyses, as only genes without alternative haplotypes are unaffected by the erroneous alignment. Here we provide a description of and protocol for read realignment (**Supplementary File**) that we hope others will find useful for generating corrected alignment files, which can then be used to generate accurate genotype calls with downstream variant calling pipelines. We have also notified the UK Biobank bioinformatics team of the bug and our proposed patch.

This study highlights the need for rigorousness and continued investigations by the community into optimal data processing protocols for UK Biobank and other large genomic resources, prompt sharing of any concerns, and timely responses to any issues raised by data guardians and providers. As tasks like sequence alignment and variant calling are very computationally expensive, robust centralized sequence data processing protocols are critical for enabling the use of such resources by the wide-ranging research community – particularly as UK Biobank prepares to expand the initial 50,000 exomes to 150,000 in early 2020, and to 500,000 whole genomes over the next few years.

## Supporting information

FE read alignment fix

## Acknowledgements

We are grateful to Dr. Olivia Osborne for helpful discussions and to William Markuske for support with high-performance computing. This work was funded by grants from the National Institute on Drug Abuse and the National Human Genome Research Institute (P50 DA037844 and R01 HG009979 to TI) as well as the National Institute for Diabetes, Digestive and Kidney diseases (K08 DK102877-01 and R03DK113328-01 to ARM) and a UCSD/UCLA Diabetes Research Center grant (P30 DK063491 to ARM).

## Declaration of Interests

TI is co-founder of Data4Cure, Inc., is on the Scientific Advisory Board, and has an equity interest. TI is on the Scientific Advisory Board of Ideaya BioSciences, Inc., has an equity interest, and receives sponsored research funding. The terms of these arrangements have been reviewed and approved by the University of California San Diego in accordance with its conflict of interest policies.

---

**Supplementary Table 1: Characteristics of 641 genes with 0 variants called in the FE pipeline exome sequences from the UK Biobank**. Variants are shown aggregated by gene in the UKB (n = 49,997) and gnomAD v2.1.1 (n = 125,748). The probability of observing at least one variant in UKB is based on the cumulative distribution function of a binomial distribution with (n = 49,997 * 2 and p = observed counts in gnomAD / (125,748 * 2)). Genes are labeled according to whether there exists an alternative contig representation in the genome reference GRCH38.

## Online Methods

### UK Biobank Whole Exome Sequencing (WES) and genotype array data

We used the sample-level aligned sequence data (CRAM files) from the Functionally Equivalent (FE) pipeline^1^. A total of 49,960 individuals have both exome sequencing data and genotype array data as of November 26, 2019, out of which 49,909 individuals pass standard genotype array quality control. As the exome data are in coordinates relative to GRCh38, but the genotype array data are in coordinates relative to GRCh37, we used the UCSC genome browser liftover tool^13^ to update genotype data coordinates to GRCh38. To facilitate direct comparison of the exome to array genotype data (**Fig. 1**), we filtered variants on the genotyping array present at a MAF > 0.01 that were also covered by the exome sequencing regions.

### Variant comparison to gnomAD

We obtained targeted exome capture regions for both UK Biobank and gnomAD^9^ (v2.1, https://storage.googleapis.com/gnomad-public/intervals/exome_calling_regions.v1.interval_list). The exome calling regions from gnomAD were converted to GRCh38 coordinates using the UCSC genome browser liftover tool^13^ to facilitate comparison to UK Biobank. We used BEDTools^14^ to extract shared regions between UKB and gnomAD. Using BEDOPS^15^, we further annotated the common genomic regions to a total of 23,040 genes based on the Ensembl 85 gene model^16^. For each gene, we aggregated variants from the UK Biobank FE pipeline project-level variant calls and compared the number of variants per gene to those in gnomAD (**Figs. 2a** and **2b**). To evaluate whether population structure contributes to the difference in variant distribution (**Fig. 2c**), we tallied the number of variants in gnomAD when subdividing individuals into six population groups: African, Latino, East Asian, European, South Asian and Other (population not assigned).

### Extraction and reprocessing of raw unmapped reads

Using SAMtools^10^, we query name sorted the aligned sequence reads in the UK Biobank CRAM files and losslessly extracted the raw unmapped reads into FASTQ files. Using BWA-MEM^12^, these reads were mapped to the full version of the GRCh38 genome reference, which contains both the primary assembly and all alternative contigs. We generated all bwa-required index files locally except the “.alt” index file, which we downloaded from the NCBI (ftp://ftp-trace.ncbi.nlm.nih.gov/1000genomes/ftp/technical/reference/GRCh38_reference_geno_me/GRCh38_full_analysis_set_plus_decoy_hla.fa.alt). We marked duplicates and recalibrated base quality scores following GATK best practices^17^. To produce the scenario in which alternative contigs are not properly referenced, we use BWA-MEM^12^ command -j to specify the aligner to ignore the “.alt” index file (**Figs. 3b**).

